# Reverting the mode of action of the mitochondrial F_O_F_1_-ATPase by *Legionella pneumophila* preserves its replication niche

**DOI:** 10.1101/2021.05.12.443790

**Authors:** Pedro Escoll, Lucien Platon, Mariatou Dramé, Tobias Sahr, Silke Schmidt, Christophe Rusniok, Carmen Buchrieser

## Abstract

*Legionella pneumophila*, the causative agent of Legionnaires’ disease, a severe pneumonia, injects *via* a type-IV-secretion-system (T4SS) more than 300 proteins into macrophages, its main host cell in humans. Certain of these proteins are implicated in reprogramming the metabolism of infected cells by reducing mitochondrial oxidative phosphorylation (OXPHOS) early after infection. Here we show that despite reduced OXPHOS, the mitochondrial membrane potential (Δ*ψ*_m_) is maintained during infection of primary human monocyte-derived macrophages (hMDMs). We reveal that *L. pneumophila* reverses the ATP-synthase activity of the mitochondrial F_O_F_1_-ATPase to ATP-hydrolase activity in a T4SS-dependent manner, which leads to a conservation of the Δ*ψ*_m_, preserves mitochondrial polarization and prevents macrophage cell death. Analyses of T4SS effectors known to target mitochondrial functions revealed that *Lp*Spl is partially involved in conserving the Δ*ψ*_m_, but not LncP and MitF. The inhibition of the *L. pneumophila-induced* “reverse mode” of the F_O_F_1_-ATPase collapsed the Δ*ψ*_m_ and caused cell death in infected cells. Single-cell analyses suggested that bacterial replication occurs preferentially in hMDMs that conserved the Δ*ψ*_m_ and showed delayed cell death. This direct manipulation of the mode of activity of the F_O_F_1_-ATPase is a newly identified feature of *L. pneumophila* allowing to delay host cell death and thereby to preserve the bacterial replication niche during infection.

## INTRODUCTION

Beyond their essential role in cellular bioenergetics, mitochondria are integrated into diverse signaling pathways in eukaryotic cells and perform various signaling functions, such as immune responses or cell death, as they play crucial roles in the regulation of apoptosis (Bock and Tait, 2020). Thus mitochondria are targeted by several intracellular bacteria during infection to modulate their functions to the bacterial advantage (Spier et al., 2019). One of these bacteria is *Legionella pneumophila*, the causative agent of Legionnaires’ disease. We have shown previously that this pathogen targets mitochondrial dynamics during infection of primary human monocyte-derived macrophages (hMDMs) by injecting type IV secretion system (T4SS) effectors such as MitF, leading to a fragmented mitochondrial network *via* the recruitment of the host fission protein DNM1L to the mitochondrial surface (Escoll et al., 2017b). Importantly, *Legionella* induced mitochondrial fragmentation at early time points such as 5 hours post-infection (hpi), when bacterial replication has not started yet, and in the absence of cell death signs. The fragmentation of mitochondrial networks provoked a T4SS-dependent reduction of mitochondrial respiration in *Legionella*-infected macrophages, evidencing a functional connection between mitochondrial dynamics and mitochondrial respiration (Escoll et al., 2017b).

Mitochondrial respiration results from coupling the activity of five complexes in the electron transport chain (ETC) at mitochondrial cristae. In this process, the reduced coenzymes NADH and FADH_2_ generated at the mitochondrial matrix by the tricarboxilic acid (TCA) cycle are oxidized at Complexes I and II where their electrons are extracted to energize the mitochondrial ETC (Nolfi-Donegan et al., 2020). The sequential transit of these electrons through Complexes I, III and IV allows to pump protons from the matrix to the intermembrane space (IMS) and at Complex IV, diatomic oxygen O_2_ serves as the terminal electron acceptor and H_2_O is formed. The increased concentration of protons [H^+^] at the IMS, compared to [H^+^] at the matrix, generates the mitochondrial membrane potential (Δ*ψ*_m_). This is necessary reveito produce ATP by fueling the rotation of Complex V, the mitochondrial F_O_F_1_-ATPase, in a process termed oxidative phosphorylation, OXPHOS (Nolfi-Donegan et al., 2020). Our previous studies determined that at 5 hpi *L. pneumophila*, by altering mitochondrial dynamics, reduced OXPHOS as well as the cellular ATP content in hMDMs in a T4SS-depended manner (Escoll et al., 2017b).

Why *L. pneumophila* and other species of intracellular bacteria reduce mitochondrial OXPHOS during infection of host cells remains a matter of debate (Escoll and Buchrieser, 2018; Russell et al., 2019). As intracellular bacteria can obtain resources only from host cells, it has been suggested that halting mitochondrial OXPHOS during infection might benefit pathogenic bacteria by redirecting cellular resources, such as glycolytic or TCA intermediates, to biosynthetic pathways that might sustain intracellular bacterial replication instead of fueling mitochondria (Escoll and Buchrieser, 2018; Russell et al., 2019). For instance, it has been shown that *Mycobacterium tuberculosis* redirects pyruvate to fatty acid synthesis and *Chlamydia trachomatis* subverts the pentose phosphate pathway to increase the synthesis of nucleotides for its own intracellular growth (Siegl et al., 2014; Singh et al., 2012). On the other hand, upon sensing bacterial lipopolysaccharides, macrophages redirect mitochondrial TCA intermediates, such as citrate or succinate, to drive specific immune functions such as the production of cytokines or the generation of antimicrobial molecules (Escoll and Buchrieser, 2019; Russell et al., 2019; O’Neill and Pearce, 2016). Thus, while these metabolic shifts, which are redirecting resources from mitochondria to the cytoplasm should be activated in macrophages to develop their antimicrobial functions, they could also benefit intracellular bacteria, as more resources would be available in the cytoplasm for bacterial growth. Importantly, reduction of OXPHOS may lead to decreased Δ*ψ*_m_ and ATP production at mitochondria, which are events that trigger the activation of cell death programs. How intracellular bacteria withdraw OXPHOS, deal with the subsequent Δ*ψ*_m_ drop and host cell death but manage to preserve their host cell to conserve their replication niche is a question that remains poorly understood.

To answer this question, we monitored the evolution of mitochondrial polarization during infection of hMDMs by *L. pneumophila*, and showed that in the absence of OXPHOS, *L. pneumophila* regulates the enzymatic activity of the mitochondrial F_O_F_1_-ATPase during infection. This allows maintaining the Δ*ψ*_m_ and delays cell death of infected hMDMs in a T4SS-dependent manner. Our results identified a new virulence mechanism of *L. pneumophila*, namely the manipulation of the mitochondrial F_O_F_1_-ATPase to preserve the integrity of infected host cells and thereby the maintenance of the bacterial replication niches.

## RESULTS

### Despite *L. pneumophila-induced* Reduction of Mitochondrial Respiration, the Mitochondrial Membrane Potential Is Maintained

We have previously shown that *L. pneumophila* strain Philadelphia JR32 impairs mitochondrial respiration during infection (Escoll et al., 2017b). Here we analyzed *L. pneumophila* strain Paris (Lpp) to learn whether this is a general characteristic of *L. pneumophila* infection. We infected hMDMs with Lpp WT or a T4SS-deficient mutant (Δ*dotA*) for 6 hours and analyzed their mitochondrial function compared to uninfected hMDMs by using a cellular *respiratory control assay* in living cells (Brand and Nicholls, 2011; Connolly et al., 2018). This assay determines oxygen consumption rate (OCR) in basal conditions and during the sequential addition of mitochondrial respiratory inhibitors. OCR variations observed indicate how mitochondrial respiration is functioning in a cell population (Figure 1A and S1A). Our results showed that basal respiration is significantly reduced (p<0. 0001) in WT-infected hMDMs compared to Δ*dotA*- and non-infected hMDMs (Figure 1A and 1B). This indicates that O_2_ consumption, which is predominantly driven by ATP turnover and the flow of H^+^ to the matrix through the mitochondrial F_O_F_1_-ATPase, is severely impaired in WT-infected hMDMs. Further analysis of OCR changes upon addition of oligomycin, an inhibitor of the mitochondrial F_O_F_1_-ATPase, indicated that the rate of mitochondrial respiration coupled to ATP synthesis is highly reduced in WT-infected hMDMs, compared to Δ*dotA*- or non-infected cells. Other respiratory parameters such as proton leak were also reduced in WT-infected macrophages (Figure 1A, S1A and S1B). Subsequent addition of an uncoupler to create a H^+^ short-circuit across the inner mitochondrial membrane (IMM), such as FCCP, allowed measuring the maximum respiration rate and the spare respiratory capacity, revealing that both were severely impaired in WT-infected cells compared to Δ*dotA*- and non-infected hMDMs (Figure 1A, S1A and S1B). Finally, inhibition of the respiratory complexes I and III with rotenone and antimycin A, respectively, measured O_2_ consumption driven by non-mitochondrial processes, such as cytoplasmic NAD(P)H oxidases, which showed similar levels of non-mitochondrial O_2_ consumption in all infection conditions (Figure 1A, S1A and S1B).

**Figure 1.**
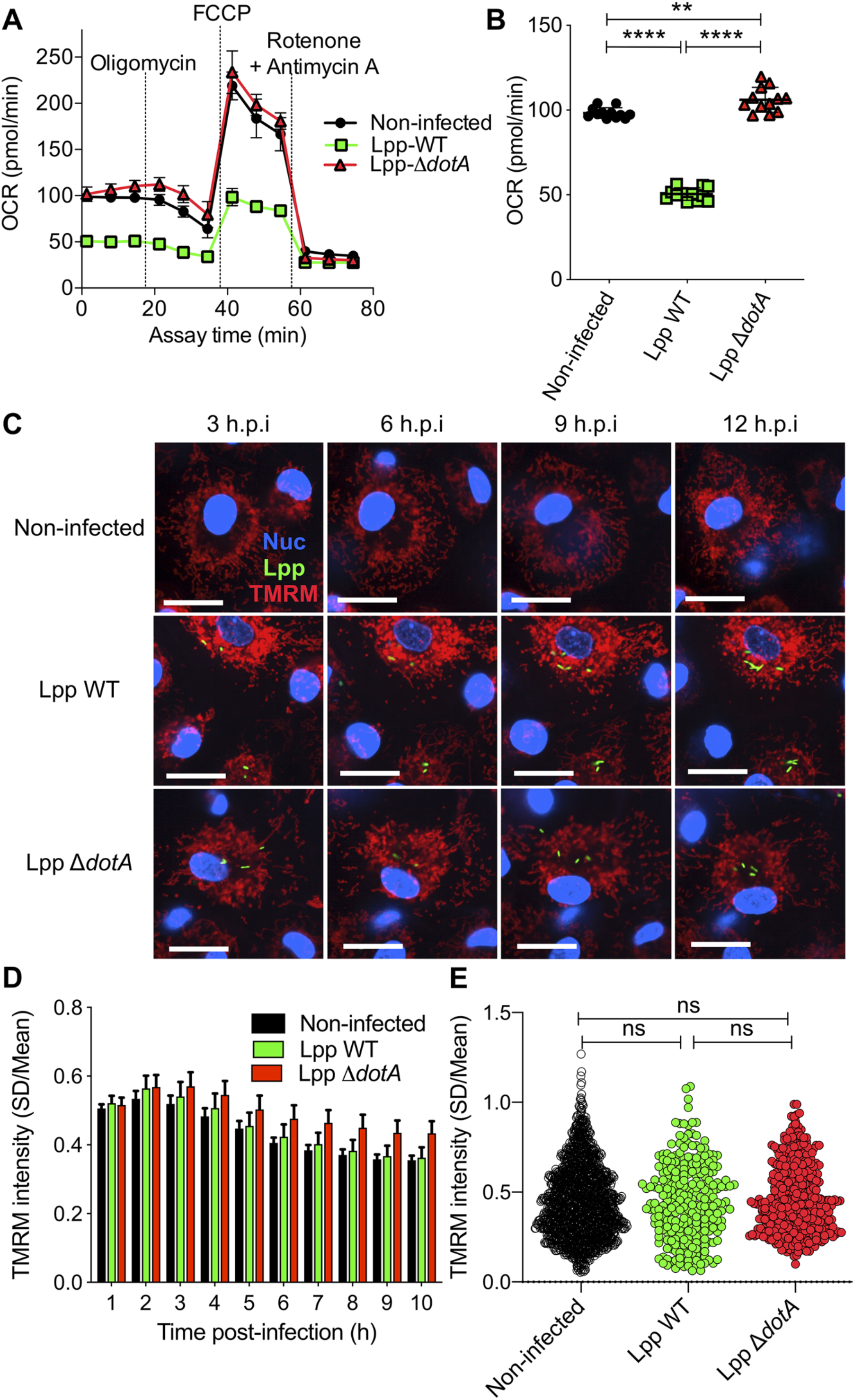
Despite a reduction of oxidative phosphorylation (OXPHOS), hMDMs maintain their Δ*ψ*_m_ during infection by *L. pneumophila*. **(A)** hMDMs were infected with *L. pneumophila* strain Paris (Lpp) wild-type (WT), a T4SS-deficient *ΔdotA* mutant, or left uninfected (Non-infected). At 6 hours post-infection (hpi), a cellular respiratory control assay was performed by measuring oxygen consumption rate (OCR) during the sequential addition of mitochondrial respiratory inhibitors (see also Figure S1A). **(B)** Basal respiration of hMDMs in the same conditions as in (A), at 6 hpi. **(C)** hMDMs were infected as in (A) with GFP-expressing bacteria (green), nuclei of host cells were stained with Hoechst (Nuc, blue) and Δ*ψ*_m_ was monitored from 1 to 12 hpi using TMRM dye in non-quenching conditions (10 nM). Representative confocal microscope images of non-infected and infected cells at 3, 6, 9 and 12 hpi are shown. Intracellular bacterial replication can be observed in Lpp-WT infected hMDMs at 12 hpi. Bar: 20 μm. **(D)** Quantification of TMRM intensity at 1-10 hpi (expressed as SD/Mean) in the assays described in (C). Data from 4 independent experiments with a total of 10 replicates **(E)** Single-cell analysis at 6 hpi of the assays described in (C). Single-cell data from a representative experiment (full time-course in Figure S1C) ** p-value < 0.01; **** p-value < 0.00001; ns = non-significant (Mann-Whitney *U* test).

Taken together, our results indicated that several mitochondrial respiration parameters were severely altered during infection with Lpp-WT, including respiration coupled to ATP production. Importantly, some of the respiratory parameters measured that are oligomycin-sensitive were reduced in Lpp-WT-infected hMDMs but not in Δ*dotA* -infected cells, suggesting that the mitochondrial F_O_F_1_-ATPase activity may be altered during *L. pneumophila* infection in a T4SS-depended manner.

The transition of electrons across mitochondrial ETC complexes allows the extrusion of H^+^from the matrix to the IMS generating a H^+^ circuit where the mitochondrial F_O_F_1_-ATPase is the dominant H^+^ re-entry site during active ATP synthesis by OXPHOS. In cellular steady-state conditions, extrusion and re-entry H^+^ fluxes across mitochondrial membranes are balanced (Brand and Nicholls, 2011). Therefore, any exogenous alteration of ATP turnover and/or F_O_F_1_-ATPase activity influences this H^+^ circuit and might be reflected in Δ*ψ*_m_ levels. Thus, we decided to quantify the Δ*ψ*_m_ in infected cells. We developed a miniaturized *high-content* assay based on kinetic measurements of TMRM fluorescence in non-quenching conditions (10nM), where TMRM fluorescence in mitochondria is proportional to the Δ*ψ*_m_ (Connolly et al., 2018; Duchen et al., 2003). This assay allowed to measure changes in the Δ*ψ*_m_ at the single-cell level and in thousands of living cells during the course of infection (Figure 1C). Image analysis showed that the Δ*ψ*_m_ slightly increased in Lpp-WT-, Lpp-Δ*dotA*- and non-infected cell populations during the first hours of infection (1-3 hpi), and progressively decreased during the time-course with no differences between the infection conditions (Figure 1D). Single-cell analyses (Figure 1E and S1C) showed that Lpp-WT-, Lpp-Δ*dotA*- and non-infected single hMDMs showed a wide range of Δ*ψ*_m_ values at any time-point (Figure S1C) with no significant differences between them at 6 hpi (Figure 1E). Thus, despite a significant reduction of OXPHOS the Δ*ψ*_m_ was maintained in infected cells, suggesting that *L. pneumophila* manipulates the mitochondrial ETC to conserve the Δ*ψ*_m_ of hMDMs in the absence of OXPHOS.

### *L. pneumophila* Infection Induces the “Reverse Mode” of the Mitochondrial F_O_F_1_-ATPase in a T4SS-depended Manner

The mitochondrial F_O_F_1_-ATPase is a fascinating molecular machine that rotates clockwise when it works in the “forward mode”, synthesizing ATP by using the Δ*ψ*_m_ generated by the H^+^ circuit (Figure 2A, left). It can also rotate counter-clockwise when it works in the “reverse mode”. In this case, it hydrolyzes ATP to maintain Δ*ψ*_m_ in the absence of OXPHOS (Figure 2A, right). As our results showed that *L. pneumophila* highly reduced OXPHOS, likely by an alteration of the FoF1-ATPase activity, while the Δ*ψ*_m_ was conserved, we investigated in which activity mode the FoF1-ATPase worked during *Legionella* infection. A widely used method to investigate the directionality of the FoF1-ATPase in intact cells is to monitor changes in Δ*ψ*_m_ after the addition of FoF1-ATPase inhibitors, such as oligomycin or DCCD (Connolly et al., 2018; Gandhi et al., 2009). These inhibitors block both modes of function, thus if the F_O_F_1_-ATPase is working in the “forward mode” the Δ*ψ*_m_ will increase after adding the inhibitor, as the inhibition of the H^+^ flux to the matrix through the ATPase leads to an accumulation of H^+^ at the IMS (Figure 2B, left). If Δ*ψ*_m_ decreases after ATPase inhibition, the F_O_F_1_-ATPase works in the “reverse mode”, since now H^+^ cannot translocate to the IMS to maintain the Δ*ψ*_m_ (Figure 2B, right). Here, we used the aforementioned TMRM high-content assay to monitor the Δ*ψ*_m_ in living hMDMs at 6 hpi, when OXPHOS is impaired and Δ*ψ*_m_ is maintained.

**Figure 2.**
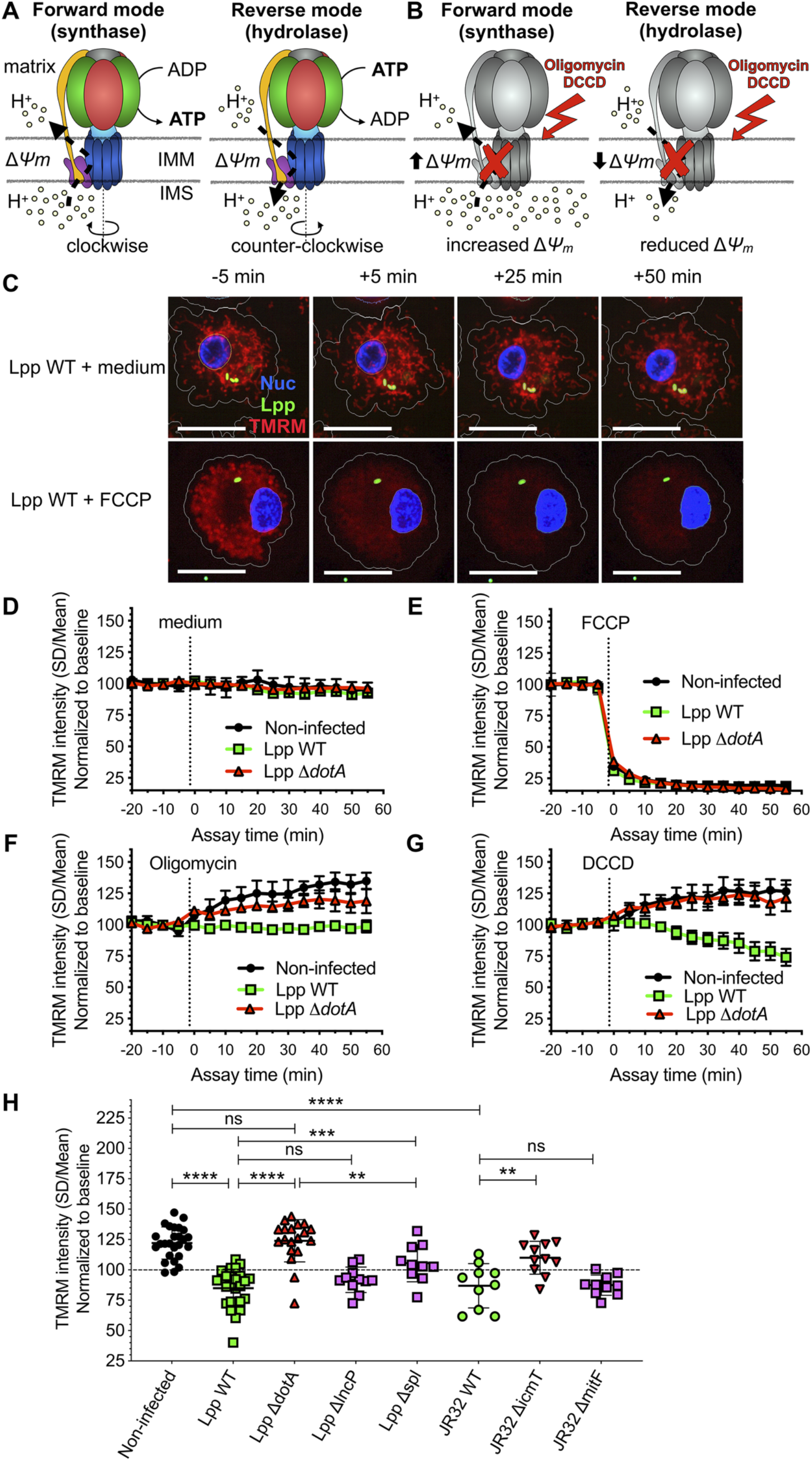
The mitochondrial F_O_F_1_-ATPase works in the “reverse mode” during infection of hMDMs by *L. pneumophila*. **(A)** In the “forward mode” of the mitochondrial ATPase, the Δ*ψ*_m_ generated by the Electron Transport Chain is used by the F_O_F_1_-ATPase to synthesize ATP. The “reverse mode” of the F_O_F_1_-ATPase leads to ATP hydrolysis to pump H^+^ to the intermembrane space (IMS). IMM: inner mitochondrial membrane. **(B)**When the F_O_F_1_-ATPase is inhibited by Oligomycin or DCCD, an increase in Δ*ψ*_m_ indicates that the ATPase was working in the “forward mode” (H^+^ accumulate in the IMS), while a decrease in Δ*ψ*_m_ indicates functioning in the “reverse mode” (H^+^ cannot be translocated to the IMS by the F_O_F_1_-ATPase to sustain the Δ*ψ*_m_). **(C)** hMDMs were infected with GFP-expressing bacteria (green), or left uninfected (Non-infected). At 5.5 hpi cells were labeled with Hoechst to identify the cell nucleus (Nuc, blue) and TMRM (red) to quantify Δ*ψ*_m_. At 6 hpi, addition of medium (no changes) or FCCP (complete depolarization) was used as controls. Representative confocal images of Lpp-WT-infected hMDMs (6 hpi) at 5 min before the addition of medium (top) or FCCP (bottom), and at 5, 25 and 50 min after addition of medium or FCCP. Bar: 20 μm. **(D)** Quantification of (C) before (baseline) and after the addition of medium. Each dot represents mean ± SD of 3 independent experiments with a total of 8 replicates. **(E)** Same as (D) but FCCP was added. **(F)** Same as (D) but oligomycin was added. **(G)** Same as (D) but DCCD was added. **(H)** Same as (C) but infection was performed with Lpp-WT, Lpp-Δ*dotA*, Lpp-Δ*lncP*, Lpp-Δ*spl*, *L. pneumophila* strain Philadelphia JR32 (JR32)-WT, JR32-Δ*icmT* or JR32-Δ*mitF*. TMRM values (SD/Mean) at 50 min after DCCD addition are shown. Data from a minimum of 3 experiments per strain with 10 or more replicates per strain ** p-value < 0.01; *** p-value < 0.001; **** p-value < 0.00001; ns = non-significant (Mann-Whitney *U* test)

First, we recorded a baseline and then added medium as a control. As expected, this did not alter Δ*ψ*_m_ in any infection condition (Figure 2C and 2D). However, the addition of FCCP completely depolarized mitochondria, leading to an abrupt drop of Δ*ψ*_m_ in Lpp-WT-, Lpp-Δ*dotA*- and non-infected hMDMs (Figure 2C and 2E), demonstrating that this assay can monitor changes in the Δ*ψ*_m_ simultaneously in hundreds of infected cells. To analyze whether the F_O_F_1_-ATPase worked in the synthase (forward) or the hydrolase (reverse) mode we added oligomycin (Figure 2F) or DCCD (Figure 2G) to the infected cells. The Δ*ψ*_m_ increased in non-infected or Δ*dotA*-infected hMDMs, which suggested that the ATPase worked in the “forward mode” in these infection conditions. In contrast, the addition of oligomycin to Lpp-WT-infected hMDMs had no effect on the Δ*ψ*_m_ (Figure 2F), while addition of DCCD decreased the Δ*ψ*_m_ (Figure 2G). Thus, our results indicate that the F_O_F_1_-ATPase worked in the “forward mode” in non-infected or Δ*dotA* -infected macrophages, whereas the F_O_F_1_-ATPase worked in the “reverse mode” during infection of hMDMs with the WT strain. This suggests that the induction of the “reverse mode” depends on the action of T4SS effector(s).

### The T4SS Effector *LpSPL* Participates in the Induction of the “Reverse Mode” of the Mitochondrial F_O_F_1_-ATPase During Infection

Among the more than 300 bacterial effectors that *L. pneumophila* injects into host cells through its T4SS (Mondino et al., 2020), three have been shown to target mitochondrial structures or functions. LncP is a T4SS effector targeted to mitochondria that assembles in the inner mitochondrial membrane (IMM) and seems to transport ATP across mitochondrial membranes (Dolezal et al., 2012). The effector *Lp*Spl (also known as LegS2) was suggested to target mitochondria (Degtyar et al., 2009), the endoplasmic reticulum (ER, (Rolando et al., 2016), and mitochondrial-associated membranes, MAMs ((Escoll et al., 2017a). *Lp*Spl encodes a sphingosine-1 phosphate (S1P) lyase that directly targets the host sphingolipid metabolism and restrains autophagy in infected cells. MitF (LegG1) activates the host small GTPase Ran to promote mitochondrial fragmentation during infection of human macrophages (Escoll et al., 2017b).

To learn if any of these effectors is involved in the T4SS-dependent induction of the “reverse mode” of the F_O_F_1_-ATPase, we infected hMDMs during 6 hours with Lpp-WT or its isogenic mutants lacking the T4SS (Lpp-Δ*dotA*), lacking the effector LncP (Lpp-Δ*lncP*), lacking the effector *Lp*Spl (Lpp-Δ*spl*), and *L. pneumophila* strain Philadelphia JR32 (JR32-WT) and its isogenic mutants lacking the T4SS (JR32-Δ*icmT*) or the effector MitF (JR32-Δ*mitF*). Using the TMRM high-content assay, we measured the Δ*ψ*_m_ after the inhibition of the F_O_F_1_-ATPase by DCCD (Figure 2H). Our results indicated that, while the F_O_-F_1_-ATPase worked in the “forward mode” in non-infected hMDMs and during infection with T4SS-deficient mutants (Lpp-Δ*dotA* and JR32-Δ*icmT*), the F_O_F_1_-ATPase worked in the “reverse mode” during infection with the Lpp-WT and JR32-WT strains (Figure 2G). Infection with Lpp-Δ*lncP* and JR32-Δ*mitF* were not significantly different compared to the WT strains, suggesting that these effectors are not involved in the induction of the “reverse mode” of the mitochondrial ATPase. However, mitochondria of cells infected with Lpp-Δ*spl* showed a significantly higher Δ*ψ*_m_ after DCCD treatment than mitochondria of cells infected with Lpp-WT (p=0.0006), and a significantly lower Δ*ψ*_m_ after DCCD treatment than mitochondria of cells infected with the Lpp-Δ*dotA* strain (p=0.0034). This suggests that *Lp*Spl is partially involved in the induction of the “reverse mode” of the F_O_F_1_-ATPase, however other additional T4SS effector(s) seem to participate in the modulation of the F_O_F_1_-ATPase activity mode.

### Inhibition of *Legionella-induced*“Reverse Mode” Collapses the Δ*ψ*_m_ And Ignites Cell Death of Infected Macrophages

To further analyze the importance of the activity mode of the F_O_F_1_-ATPase during infection, we used BTB06584 (hereafter called BTB), a specific inhibitor of the “reverse mode” of the mitochondrial F_O_F_1_-ATPase (Ivanes et al., 2014). We used the TMRM high-content assay and added BTB to non-infected, Lpp-WT- or Lpp-Δ*dotA*-infected hMDMs at 6 hpi. As shown in Figure 3A, the Δ*ψ*_m_ collapsed specifically and significantly in WT infected cells (Figure 3B and 3C) compared to non-infected (p=0.0022) and Δ*dotA*-infected cells (p=0.0238), further confirming that the F_O_F_1_-ATPase works in the “reverse mode” during WT infection. Indeed, the addition of BTB to Lpp-WT-infected hMDMs led to a significant reduction of the Δ*ψ*_m_ (p<0. 0001) at every time point post-infection (1-10 hpi) and at the single-cell level compared to non-treated Lpp-WT-infected cells (Figure 3D), further confirming that conservation of Δ*ψ*_m_ during *L. pneumophila* infection is caused by induction of Fo-F1-ATPase “reverse mode”.

**Figure 3.**
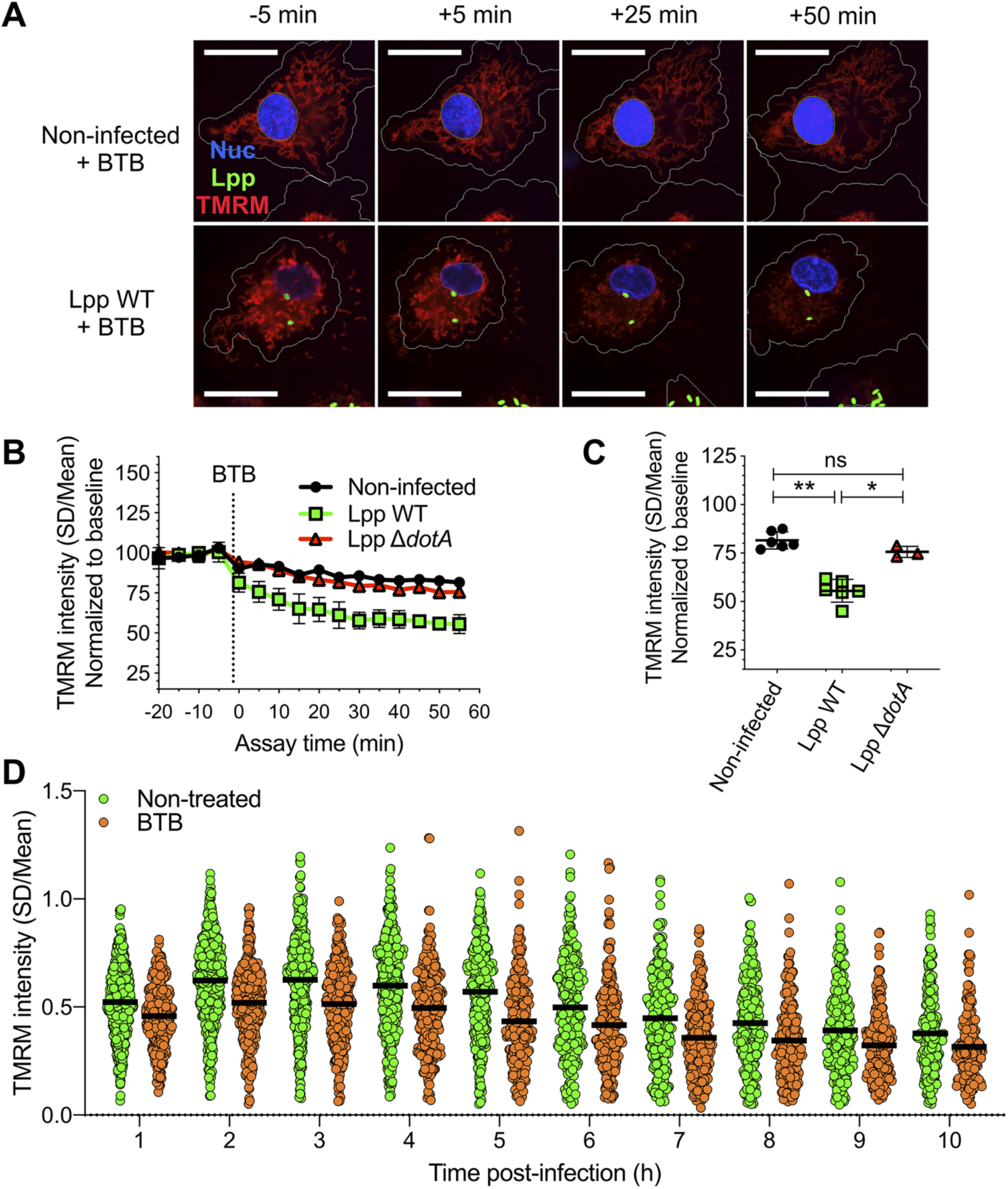
Inhibition of the “reverse mode” of mitochondrial F_O_F_1_ ATPase reduces the Δ*ψ*_m_ of *L. pneumophila-infected* hMDMs. **(A)** hMDMs were infected with GFP-expressing bacteria (green), Lpp-WT or Lpp-Δ*dotA*, or left uninfected (Non-infected). At 5.5 hpi cells were labeled with Hoechst to identify cell nucleus (Nuc, blue) and TMRM (red) to quantify Δ*ψ*_m_. At 6 hpi, BTB06584 (BTB, 50 μM), a specific inhibitor of the “reverse mode” of the ATPase, was added and Δ*ψ*_m_ monitored. Representative confocal microscopy images of non-infected (top) and Lpp-WT-infected (bottom) hMDMs (6 hpi) at 5 min before the addition and at 5, 25 and 50 min after the addition of BTB. Bar: 20 μm. **(B)** Quantification of (C) before (baseline) and after the addition of BTB. Each dot represents the mean ± SD of 3 independent experiments with a total of 6 replicates. **(C)** Same infection conditions than (A) but TMRM values (SD/Mean) at 50 min after BTB addition are shown. Data from 3 experiments with a total of 6 replicates (3 replicates for Lpp-Δ*dotA*) **(D)** Single-cell analysis of Δ*ψ*_m_ in Lpp-WT-infected hMDMs treated with BTB (50 μM) or left untreated (non-treated). Single-cell data from one representative experiment * p-value < 0.05; ** p-value < 0.01; ns = non-significant (Mann-Whitney *U* test)

As OXPHOS cessation and Δ*ψ*_m_ collapse can trigger cell death, we reasoned that induction of the “reverse mode” of mitochondrial ATPase by *L. pneumophila* to maintain Δ*ψ*_m_ in the absence of OXPHOS might delay cell death of infected cells. To test this hypothesis, we infected hMDMs with Lpp-WT and treated them with BTB or left them untreated, and then measured the percentage of living cells among infected cells (Figure 4A). Our results showed that the percentage of living, infected cells significantly decreased after 10 hpi in BTB-treated infected hMDMs compared to non-treated cells. As this reduction in the percentage of living, infected cells upon “reverse mode” inhibition might be caused by increased cell death, we used our high-content assay to measure Annexin-V, a marker of early apoptosis, in a high number of living hMDMs during infection (Figure 4B and 4C). While addition of BTB during 24 hours did not increase the percentage of Annexin-V^+^ cells in non-infected cells, the addition of this “reverse mode” inhibitor to Lpp-WT-infected hMDMs significantly increased the percentage of Annexin-V^+^ cells compared to non-treated cells (p=0.0312). This suggests that the inhibition of the “reverse mode” by BTB leads to a reduction in the percentage of infected cells as increased cell death occurs specifically in infected cells. Single-cell analysis at 12 and 18 hpi (Figure 4D and S2A) also showed higher levels of Annexin-V intensity in BTB-treated Lpp-WT-infected hMDMs compared to non-treated infected cells (p<0. 0001). BTB-treated infected cells also showed higher Hoechst nuclear levels compared to non-treated infected cells, a sign of nuclear condensation typical of apoptotic cells (Figure S2B), which further indicates that inhibition of the *Legionella*-induced ATPase “reverse mode” ignites cell death of infected macrophages. Taken together, our results suggest that the *Legionella*-induced “reverse mode” of the mitochondrial F_O_F_1_-ATPase aids to conserve Δ*ψ*_m_ during infection to delay cell death of infected macrophages.

**Figure 4.**
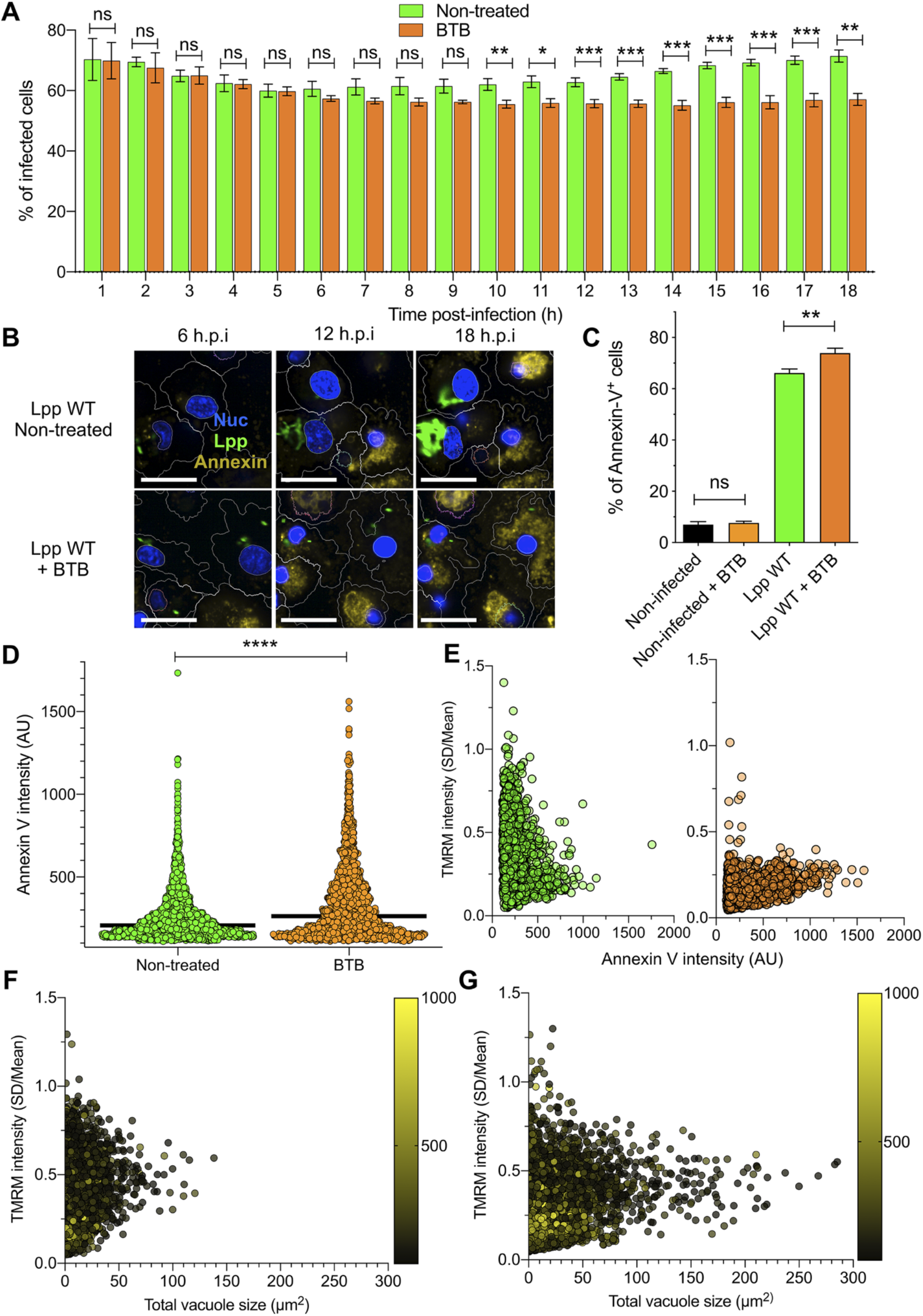
Inhibition of F_O_-F_1_ ATPase “reverse mode” increases cell death in *L. pneumophila-infected* hMDMs. **(A)** hMDMs were infected with Lpp-WT-GFP and were non-treated or treated with 50 μM BTB. The presence of GFP-expressing bacteria in each cell was monitored and the number of infected cells in the whole population was graphed as percentage of infected cells. Data from 3 independent experiments with a total of 7 replicates per condition and time-point **(B)** hMDMs were infected with Lpp-WT-GFP (green), the nuclei of host cells were stained with Hoechst (Nuc, blue) and Annexin-V Alexa Fluor 647 was added to the cell culture to monitor early cell death (Annexin, yellow) from 1 to 18 hpi in non-treated or BTB-treated hMDMs. Representative confocal images of non-treated and Lpp-WT-GFP-infected cells at 6, 12 and 18 hpi are shown. Intracellular bacterial replication can be observed in non-treated Lpp-WT infected hMDMs at 12 and 18 hpi. Bar: 20 μm.**(C)** hMDMs stained as in (B) were infected with Lpp-WT-GFP or left uninfected (Non-infected), and then were treated or not with BTB (50 μM). Percentage of Annexin-V^+^ cells at 24 hpi is shown. Data from 3 independent experiments with a total of 7 replicates per condition **(D)** Single-cell analysis (12 hpi) of Annexin-V intensity of the assays described in (B). Single-cell data from one representative experiment (18 hpi shown in Figure S2A) **(E)** hMDMs were infected with Lpp-WT-GFP, nuclei of host cells were stained with Hoechst, and TMRM and Annexin-V Alexa Fluor 647 were added to the cells to simultaneously monitor (1-18 hpi) Δ*ψ*_m_ and early cell death, respectively, in non-treated or BTB-treated hMDMs (representative multi-field confocal images in Figure S2C). Single-cell analyses (12 hpi) of Δ*ψ*_m_ (TMRM SD/Mean) and cell death (Annexin-V intensity) in more than 1600 cells per condition are shown. Single-cell data from one representative experiment; Green dots: Non-treated Lpp-WT-infected single cells. Orange dots: BTB-treated Lpp-WT-infected single cells. **(F)** Same infection conditions as in (E) but vacuole size was monitored in each Lpp-WT-infected single cell. Single-cell analyses (12 hpi) of Δ*ψ*_m_ (TMRM SD/Mean), vacuole size (μm^2^), and cell death (Annexin-V intensity) in more than 3800 cells are shown. Single-cell data from one representative experiment; Color scale (yellow) represents Annexin V intensity per cell (AU). **(G)** Same as in (F) at 18 hpi. * p-value < 0.05; ** p-value < 0.01; *** p-value < 0.001; **** p-value < 0.00001; ns = non-significant (Mann-Whitney *U* test)

To simultaneously monitor Δ*ψ*_m_ and early cell death signs in the same infected cell, we multiplexed Annexin-V and TMRM signals in our living cell assay (Figure 4E, S2C and S2D). *L. pneumophila-infected* macrophages where the “reverse mode” activity of the F_O_F_1_-ATPase was inhibited by BTB suffered a collapsed Δ*ψ*_m_ and showed higher levels of Annexin-V at 12 and 18 hpi (Figure 4E and S2D) compared to non-treated cells. Thus both events, collapse of the Δ*ψ*_m_ and triggered cell death, occurred in the same infected cell when the *Legionella*-induced “reverse mode” activity of the F_O_F_1_-ATPase was inhibited. Furthermore, when the size of the bacterial vacuole was correlated with the Δ*ψ*_m_ and cell death at the single-cell level (12 and 18 hpi, Figure 4F and 4G respectively), intracellular bacterial replication occurred preferentially in those infected macrophages with intermediate levels of TMRM (non-collapsed Δ*ψ*_m_) and low Annexin-V levels (yellow color scale, Figure 4F and 4G), further indicating that conservation of Δ*ψ*_m_ and delayed cell death are needed to guarantee the survival of infected macrophages to ensure bacterial replication.

## DISCUSSION

We show here that by inducing the “reverse mode” of the mitochondrial F_O_F_1_-ATPase, *L. pneumophila* circumvents the collapse of Δ*ψ*_m_ and cell death caused by OXPHOS cessation in infected cells. This mechanism, which partially involves the T4SS effector *Lp*Spl, maintains the Δ*ψ*_m_ and delays host cell death during infection, thus preserving bacterial replication niches in conditions where mitochondrial respiration is abruptly reduced. Indeed, not only *L. pneumophila*, but also several other intracellular bacterial pathogens, such as *Mycobacterium tuberculosis* or *Chlamydia pneumoniae*, reduce mitochondrial OXPHOS during infection (Siegl et al., 2014; Singh et al., 2012)(Escoll and Buchrieser, 2019). OXPHOS reduction allows the pathogen to redirect cellular resources from mitochondria to the cytoplasm, which enhances glycolysis and biosynthetic pathways that can provide intracellular bacteria with resources needed for bacterial growth (Escoll and Buchrieser, 2018; Russell et al., 2019). In contrast, OXPHOS cessation in macrophages also enhances the biosynthetic pathways leading to the synthesis of cytokines and antimicrobial compounds (O’Neill and Pearce, 2016; Russell et al., 2019). Furthermore, OXPHOS reduction may trigger profound consequences for host cells, such as the collapse of Δ*ψ*_m_ that may lead to subsequent cell death of infected cells. For macrophages, cell death is considered as a defense mechanism against infection (Chow et al., 2016). Indeed, pyroptosis of infected macrophages permits the spread of inflammatory mediators such as IL-1β. Thus, for intracellular bacteria, many of which infect macrophages (Mitchell et al., 2016), the death of their host cell is an obstacle as their cellular replication niche is destroyed. Therefore, while bacterial-induced reduction of OXPHOS might be beneficial for intracellular bacteria to obtain host cell resources, they need to counterbalance the consequences of OXPHOS cessation, i.e. the collapse of the mitochondrial Δ*ψ*_m_ and subsequent cell death, to preserve their replication niches.

We have previously shown that the *L. pneumophila* T4SS effector MitF is implicated in fragmenting the mitochondrial networks of infected macrophages. These changes in the mitochondrial dynamics have a profound impact on OXPHOS that was severely impaired and accompanied by increased glycolysis in *Legionella-infected* cells (Escoll et al., 2017b). Here we show that, despite the impairment of mitochondrial respiration in infected cells, *L. pneumophila* conserves the Δ*ψ*_m_ of host cells by inducing the “reverse mode” of the F_O_F_1_-ATPase by a mechanism that is T4SS-dependent and partially mediated by the T4SS effector *Lp*Spl. When translocated into human cells, the S1P-lyase activity of *L. pneumophila Lp*Spl reduces S1P levels in infected cells and restrains autophagy, likely because S1P is involved in the initiation of autophagosome formation at MAMs (Rolando et al., 2016).

How *Lp*Spl may regulate the activity of the F_O_F_1_-ATPase is an interesting question. Indeed, phosphorylated lipids are critical regulators of mitochondrial functions and S1P is a potent lipid mediator that regulates various physiological processes as well as diverse mitochondrial functions such as mitochondrial respiration, ETC functioning or mitochondrial-dependent cell death (Hernández-Corbacho et al., 2017; Nielson and Rutter, 2018). Furthermore it was reported that S1P interaction with Prohibitin 2 (PHB2) regulates ETC functioning and mitochondrial respiration (Strub et al., 2011) and that a link between PHB2, ETC functioning and the activation of “mitoflashes” (Jian et al., 2017) exists, which are dynamic and transient uncouplings of mitochondrial respiration from ATP production that are partially dependent on the “reverse mode” of the F_O_F_1_-ATPase (Wei-LaPierre and Dirksen, 2019). Thus, it is possible that *Lp*Spl modulates mitochondrial S1P levels helping the induction of the “reverse mode” of the mitochondrial F_O_F_1_-ATPase by involving PHB2, ETC complex assembly or the generation of mitoflashes, a fascinating possibility that we will further investigate.

The regulation of host cell death by intracellular bacteria is widely studied (Rudel et al., 2010). For *L. pneumophila*, T4SS effectors activating and inhibiting cell death of infected cells have been described (Speir et al., 2014), suggesting that a very delicate interplay of positive and negative signals governs the fate of infected macrophages. Here we have shown that bacterial replication occurs preferentially in those infected macrophages that are able to conserve the Δ*ψ*_m_ and delay cell death, a condition that is difficult to achieve in the absence of mitochondrial respiration. Thus, the manipulation of the activity of the mitochondrial F_O_F_1_-ATPase by *L. pneumophila*, which allows this pathogen to use the ATP hydrolase activity to pump H^+^ to the IMS to maintain the Δ*ψ*_m_ in infected cells, is a novel virulence strategy that might contribute to the fine-tuning of the timing of host cell death during bacterial infection.

## MATERIALS and METHODS

### Human Primary Cell Cultures

Human blood was collected from healthy volunteers under the ethical rules established by the French National Blood Service (EFS). Peripheral blood mononuclear cells (PBMCs) were isolated by Ficoll-Hypaque density-gradient separation (Lympholyte-H; Cedarlane Laboratories) at room temperature. PBMCs were incubated with anti-human CD14 antibodies coupled to magnetic beads (Miltenyi Biotec) and subjected to magnetic separation using LS columns (Miltenyi Biotec). Positive selected CD14^+^ cells were counted and CD14 expression was analysed by flow cytometry, repeatedly showing a purity > 90%. CD14 cells were plated in RPMI 1640 medium (Life Technologies) supplemented with 10% heat-inactivated fetal bovine serum (FBS, Biowest) in 6 well multi-dish Nunc UpCell Surface cell culture plates or 10 cm Nunc UpCell Surface cell culture dishes (Thermo Fisher) and differentiated to human monocyte-derived macrophages (hMDMs) by incubation with 100 ng/ml of recombinant human macrophage colony-stimulating factor (rhMCSF, Miltenyi Biotec) for 6 days at 37°C with 5% CO_2_ in a humidified atmosphere. At day 3, additional rhMCSF (50 ng/ml) was added. After 6 days differentiation, UpCell plates were placed at 20°C during 10 minutes and hMDMs were gently detached, counted and plated in RPMI 1640 10% FBS in 384-well plates (Greiner Bio-One).

### Bacterial strains and mutant construction

*L. pneumophila* strain Paris or JR32 and their derivatives were grown for 3 days on N-(2-acetamido)-2-amino-ethanesulfonic acid (ACES)-buffered charcoal-yeast (BCYE) extract agar, at 37°C. For eGFP-expressing strains harbouring pNT28 plasmid (Tiaden et al., 2007), chloramphenicol (Cam; 5 μg/mL) was added. Knock-out mutant strains of *L. pneumophila* genes coding for the T4SS effectors *Lp*Spl and MitF/LegG1 were previously described (Escoll et al., 2017b; Rolando et al., 2016; Rothmeier et al., 2013). The knock-out mutant strain of the *L. pneumophila* gene coding for the effector LncP was constructed as previously described (Brüggemann et al., 2006; Rolando et al., 2016). In brief, the gene of interest was inactivated by introduction of an apramycine resistance (apraR) cassette into the chromosomal gene by 3-steps PCR. The g primers used for the *lncP* (lpp2981) knock out mutant are: LncP_F: ACCCTGGTTCATGGTAACAATGG; LncP_Inv_R: GAGCGGATCGGGGATTGTCTTATCAGGCGAATGGTGTGAAAGG; LncP_Inv_F: GCTGATGGAGCTGCACATGAAACGTCATGGTCGTGCTGGTTG; LncP_R: AATCAGATGGGTAAGCCGATTGG. To amplify the apramycine cassette, the primers Apra_F: TTCATGTGCAGCTCCATCAGC and Apra_R: AAGACAATCCCCGATCCGCTC were used.

### Infection of hMDMs and automatic confocal imaging

hMDMs were infected with *L. pneumophila* grown for three days on BCYE agar plates. Bacteria were dissolved in 1X PBS (Life Technologies), the optical density (OD) was adjusted to OD_600_ of 2.5 (2.2 × 10^9^ bacteria/mL) and the bacteria were then further diluted in serum-free XVIVO-15 medium (Lonza) prior to infection to obtain the respective multiplicity of infection (MOI). hMDMs were washed twice with serum-free XVIVO-15 medium and then infected (MOI = 10) with 25 μL of bacteria in 384-well plates (Greiner Bio-One). The infection was synchronized by centrifugation (200 g for 5 min) and the infected cells were incubated at 37°C for 5 min in a water bath and then for 25 min at 37°C/5%CO_2_. After three intensive washes with serum-free XVIVO-15 medium, the infection proceeded in serum-free XVIVO-15 medium for the respective time points. 30 min prior imaging, 25 μL of culture medium were removed and replaced by 25 μL of 2X mix of dyes, to a final concentration of 200 ng/mL of Hoechst H33342 (nuclear staining; Life Technologies), 10 nM of TMRM (mitochondrial membrane potential; Life Technologies), and/or 1/100 Annexin-V-Alexa Fluor 647 (early apoptosis; Life Technologies). If chemical inhibitors of the Electron Transport Chain (ETC) were used in the experiments, they were added to hMDMs at the indicated times points at the following concentrations: 5 μM Oligomycin (Enzo), 100 μM DCCD (Dicyclohexylcarbodiimide, Sigma), 10 μM FCCP (Tocris), 50 μM BTB06584 (Sigma). Image acquisitions of multiple fields (9 to 25) per well were performed on an automated confocal microscope (OPERA Phenix, Perkin Elmer) using 60X objective, excitation lasers at 405, 488, 561 and 640 nm, and emission filters at 450, 540, 600 and 690 nm, respectively.

### Metabolic Extracellular Flux Analysis

hMDMs (50,000) were plated in XF-96-cell culture plates (Seahorse Bioscience). For OCR measurements, XF Assay Medium (Seahorse Bioscience) supplemented with 1 mM pyruvate and 10 mM glucose was used, and OCR was measured in a XF-96 Flux Analyzer (Seahorse Bioscience). For the mitochondrial respiratory control assay, hMDMs were infected at MOI = 10 and at 6 hpi., different drugs were injected (Mitostress kit, Seahorse Bioscience) while OCR was monitored. Specifically, Olygomycin was injected through the port A, then FCCP was injected through the port B, and finally Rotenone + Antimycin A were injected through the port C, to reach each of the drugs a final concentration in the well of 0.5 μM.

### Automatic High-Content Analyses (HCA)

All analyses were performed with Harmony software v.4.9 (Perkin Elmer) using in-house developed scripts (available upon request). For the HCA of the mitochondrial membrane potential (Δ*ψ*_m_), the Hoechst signal was used to segment nuclei in the 405/450 channel (excitation/emission), Hoechst background signal in the cytoplasm was used to segment the cytoplasm region in the 405/450 channel, *L. pneumophila* was identified by measuring the GFP signal in the 488/540 channel, and TMRM (10 nM) signal in the 561/600 channel was used to measure Δ*ψ*_m_ by calculating SD/Mean TMRM intensity values in each infected and non-infected cell. For the HCA of cell death, the Hoechst signal was used to segment nuclei in the 405/450 channel, Hoechst background signal in the cytoplasm was used to segment the cytoplasm region, and the identification of *L. pneumophila* was performed using the GFP signal in the 488/540 channel. Then, Annexin-V-AlexaFluor 647 signal was measured in the 640/690 channel and the Hoechst signal intensity was measured in the 405/450 channel for each infected or non-infected cell. For the HCA analyses combining Δ*ψ*_m_ and cell death, both HCA strategies aforementioned were merged, using high Hoechst signal in the 405/450 channel to segment nuclei, low Hoechst signal in the 405/450 channel to segment cytoplasm, GFP signal in the 488/540 channel to identify bacteria, TMRM signal in the 561/600 channel to measure Δ*ψ*_m_ (SD/Mean) and Annexin-V-AlexaFluor 647 signal in the 640/690 channel to measure cell death.

### Whole genome sequencing for mutant validation

Chromosomal DNA was extracted from BCYE-grown *L. pneumophila* using the DNeasy Blood and Tissue Kit (Qiagen). The Illumina NGS libraries were prepared using the Nextera DNA Flex Library Prep following the manufacturer’s instructions (Illumina Inc.). High-throughput sequencing was performed with a MiSeq Illumina sequencer (2 × 300 bp, Illumina Inc.) by the Biomics Pole (Institut Pasteur). For the analysis, we first removed adapters from Illumina sequencing reads using Cutadapt software version 1.15 (Martin, 2011) and we used Sickle (https://github.com/najoshi/sickle) with a quality threshold of 20 (Phred score) to trim bad quality extremities. Reads were assembled using Spades (Nurk et al., 2013) and different K-mer values. The region corresponding to the gene of interest was identified by blastn, extracted, and compared to the homologous region in the *L. pneumophila* strain Paris WT genome and to the antibiotic cassette sequence using blastn. The results are visually inspected with ACT (Artemis Comparison Tool) (Carver et al., 2005). In addition, we searched the entire genome whether off-target mutations had occurred, using Bowtie 2 (Langmead and Salzberg, 2012) to perform a mapping against the genome sequence of *L. pneumophila* strain Paris (NC_006368.1). SNPs and small indels were searched for with freebayes SNP caller (Garrison and Marth, 2012), mutations and small indels were visualized in the Artemis genome viewer (Carver et al., 2005) to analyze them (new amino acid, synonymous mutation, frameshifts, etc). We used Samtools to find regions with no coverage (or close to zero) (Li et al., 2009). Regions or positions with such anomalies were visualized and compared with the corresponding region of the assembly. This confirmed that no off-target mutations impacting the phenotype of the mutant had occurred.

### Statistical analyses

The two-sample Student’s t-test (Mann-Whitney *U* test, non-assumption of Gaussian distributions) was used in all data sets unless stated otherwise. Data analysis was performed using Prism v9 (Graphpad Software).

## AUTHOR CONTRIBUTIONS

PE conceived the study; PE, LP, MD and SS prepared blood-derived human cells and/or performed experiments; TS constructed bacterial mutants; PE, LP and CR analyzed data; PE and CB provided funding; CB provided critical advice; PE and CB wrote the manuscript.

## ACKNOWLEDGEMENTS

We acknowledge C.B.’s, P. Glaser’s lab members and F. Stavru at Institut Pasteur for fruitful discussions. We thank N. Aulner, A. Danckaert, Photonic BioImaging (PBI) UTechS and M. Hassan, Center for Translational Science (CRT) at Institut Pasteur, for support. This research was funded by the Institut Pasteur, DARRI - Institut Carnot - *Microbe et santé* (grant number INNOV-SP10-19) to PE; the Agence National de Recherche (grant number ANR-10-LABX- 62-IBEID) and the Fondation de la Recherché Médicale (FRM) (grant number EQU201903007847) to CB, and the Région Ile-de-France (program DIM1Health) to PBI (part of FranceBioImaging, ANR-10-INSB-04-01). MD was supported by the Ecole Doctorale FIRE – “Programme Bettencourt”. SS was supported by the Pasteur Paris-University (PPU) International PhD Program.

