## Supplemental Material for "Reverting the mode of action of the mitochondrial F_O_F_1_-ATPase by *Legionella pneumophila* preserves its replication niche"

**Figure S1: Bioenergetic profile during infection by *L. pneumophila*.**

**Figure S2: Inhibition of $F_O$-$F_1$ ATPase "reverse mode" increases cell death in *L. pneumophila*-infected hMDMs.**

### SUPPLEMENTARY FIGURE LEGENDS

**Figure S1. Bioenergetic profile during infection by *L. pneumophila*. (A)** Bioenergetic profiles of the key parameters of mitochondrial respiration during a mitochondrial respiratory control assay using the Seahorse XF Mitostress kit. Sequential compound injections measure basal respiration, ATP production, proton leak, maximal respiration, spare respiratory capacity, and non-mitochondrial respiration (Source: Seahorse Bioscience). **(B)** hMDMs were infected with *L. pneumophila* strain Paris (Lpp) wild-type (WT), a T4SS-deficient Δ*dotA* mutant, or left uninfected (Non-infected). At 6 hours post-infection (hpi), a cellular respiratory control assay (Seahorse) was performed by measuring oxygen consumption rate (OCR) during the sequential addition of mitochondrial respiratory inhibitors. Quantification of key parameters of mitochondrial functioning, derived from this assay (Figure 1A), are shown. **(C)** hMDMs were infected as in (B) with GFP-expressing bacteria, nuclei of host cells were stained with Hoechst and $\Delta\psi_m$ was monitored using TMRM dye in non-quenching conditions (10 nM). Single-cell analysis of TMRM intensity at 1-10 hpi (expressed as SD/Mean) is shown.

**Figure S2. Inhibition of $F_O$-$F_1$ ATPase "reverse mode" increases cell death in *L. pneumophila*-infected hMDMs. (A)** hMDMs were infected with Lpp-WT-GFP, the nuclei of host cells were stained with Hoechst and Annexin-V-647 was added to the cell culture to monitor early cell death from 1 to 18 hpi in non-treated or BTB-treated hMDMs. Single-cell analysis of Annexin-V intensity at 18 hpi is shown. **(B)** hMDMs were infected as in (A) and Hoechst intensity in the nucleus was analyzed in single cells at 12 hpi. (C) hMDMs were infected with Lpp-WT-GFP (green), nuclei of host cells were stained with Hoechst (Nuc, blue), and TMRM (red) and Annexin-V Alexa Fluor 647 (yellow) were added to the cells to simultaneously monitor $\Delta\psi_m$ and early cell death, respectively, in non-treated or BTB-treated hMDMs, respectively. Confocal images of living infected cells were automatically acquired in 16 fields per well (4 wells per condition) using 60X magnification. Representative images of 16 stitched fields per condition are shown at 6, 12 and 18 hpi. Bar: 200 µm. **(D)** hMDMs were infected with Lpp-WT-GFP, nuclei of host cells were stained with Hoechst, and TMRM and Annexin-V Alexa Fluor 647 were added to the cells to simultaneously monitor (1-18 hpi) $\Delta\psi_m$ and early cell death, respectively, in non-treated or BTB-treated hMDMs, respectively. Single-cell analyses (18 hpi) of $\Delta\psi_m$ (TMRM SD/Mean) and cell death (Annexin-V intensity) in more than 1600 cells per condition are shown. Single-cell data from one representative experiment; Green dots: Non-treated Lpp-WT-infected single cells. Orange dots: BTB-treated Lpp-WT-infected single cells.

Figure S1

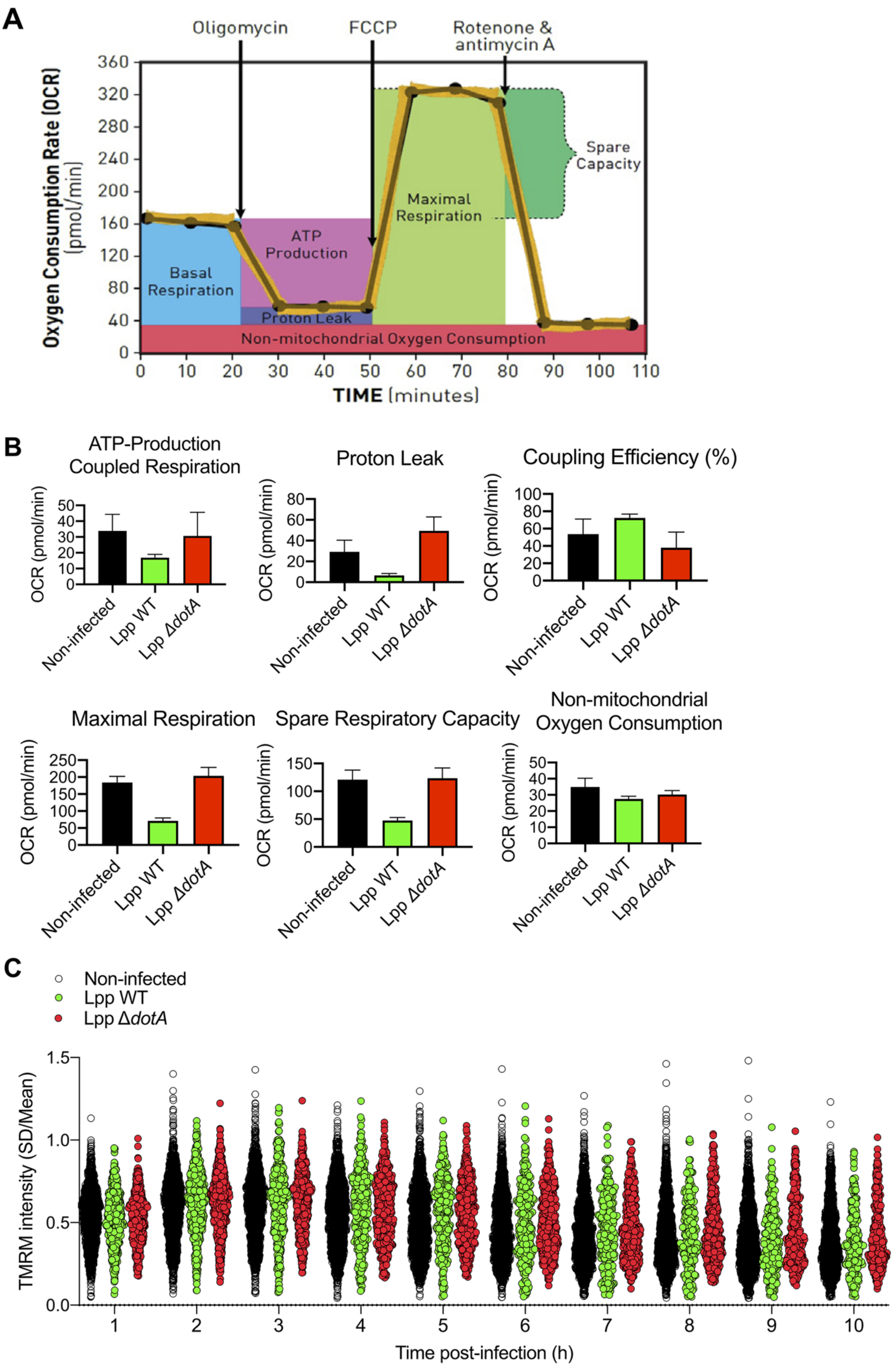
A
Oligomycin
FCCP
Rotenone & antimycin A
Oxygen Consumption Rate (OCR) (pmol/min)
Spare Capacity
Maximal Respiration
ATP Production
Basal Respiration
Proton Leak
Non-mitochondrial Oxygen Consumption
TIME (minutes)
B
ATP-Production Coupled Respiration
Proton Leak
Coupling Efficiency (%)
Maximal Respiration
Spare Respiratory Capacity
Non-mitochondrial Oxygen Consumption
OCR (pmol/min)
Non-infected
Lpp WT
Lpp ΔdotA
C
Non-infected
Lpp WT
Lpp ΔdotA
TMRM intensity (SD/Mean)
Time post-infection (h)

Figure S2

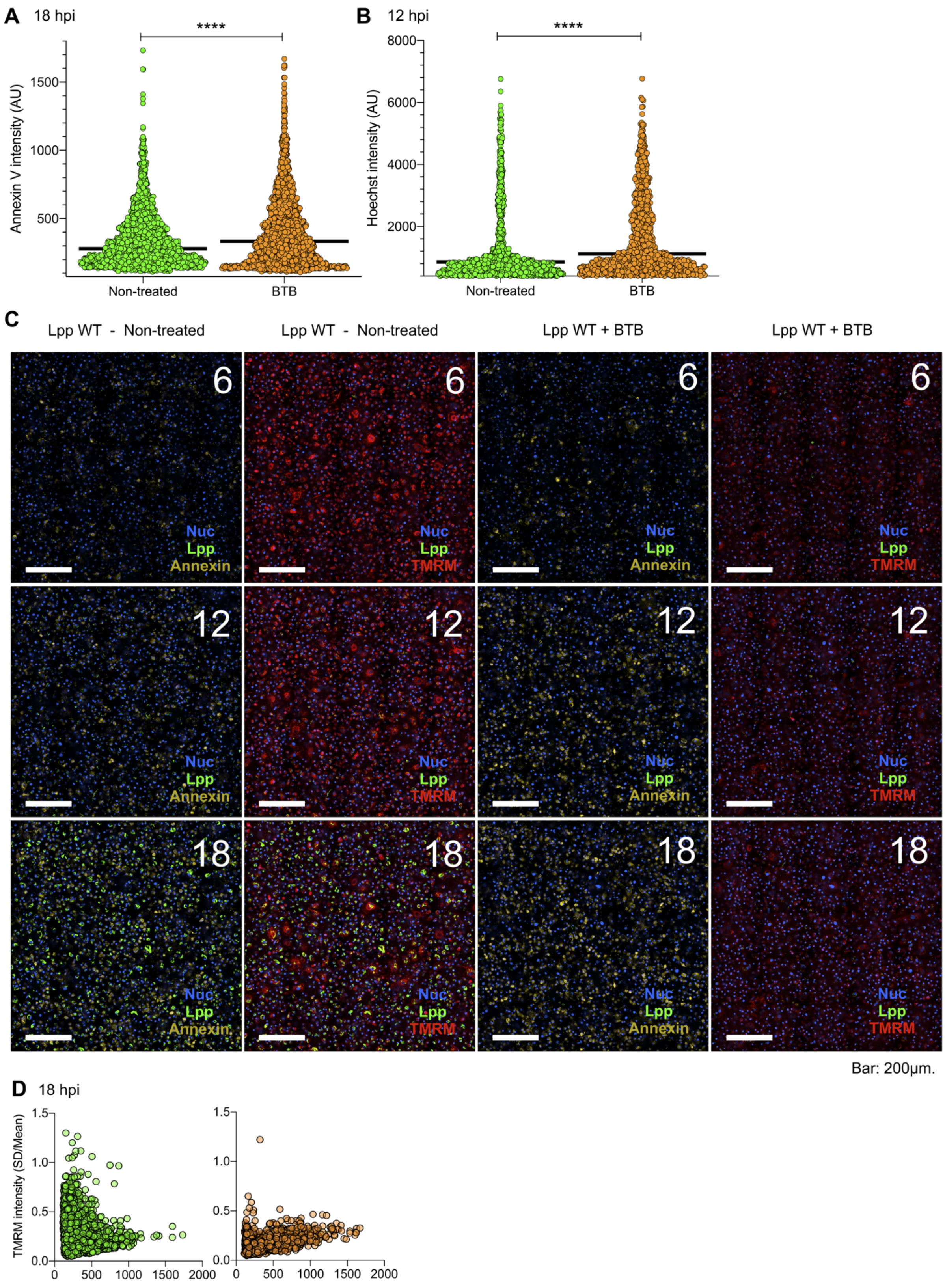
A 18 hpi
****
Annexin V intensity (AU)
1500
1000
500
Non-treated
BTB
B 12 hpi
****
Hoechst intensity (AU)
8000
6000
4000
2000
Non-treated
BTB
C
Lpp WT - Non-treated
Lpp WT - Non-treated
Lpp WT + BTB
Lpp WT + BTB
6
12
18
Nuc
Lpp
Annexin
TMRM
Bar: 200μm.
D 18 hpi
TMRM intensity (SD/Mean)
1.5
1.0
0.5
0.0
0
500
1000
1500
2000
